# Mitochondrial ATP production is required for endothelial cell control of vascular tone

**DOI:** 10.1101/2021.09.14.460297

**Authors:** Calum Wilson, Matthew D. Lee, Charlotte Buckley, Xun Zhang, John G. McCarron

## Abstract

Arteries and veins are lined by non-proliferating endothelial cells that play a critical role in regulating blood flow. Endothelial cells also regulate tissue perfusion, metabolite exchange, and thrombosis. It is thought that endothelial cells rely on ATP generated via glycolysis to fuel each of these energy-demanding processes. However, endothelial metabolism has mainly been studied in the context of proliferative cells in angiogenesis, and little is known about energy production in endothelial cells within the fully-formed vascular wall. Using intact arteries isolated from rats and mice, we show that inhibiting mitochondrial oxidative phosphorylation disrupts endothelial control of vascular tone. The role for endothelial cell energy production is independent of species, sex, or vascular bed. Basal, mechanically-activated, and agonist-evoked calcium activity in intact artery endothelial cells are each prevented by inhibiting mitochondrial ATP synthesis. This effect is mimicked by blocking the transport of pyruvate, the master fuel for mitochondrial energy production, through the mitochondrial pyruvate carrier. These data show that mitochondrial ATP is necessary for calcium-dependent, nitric oxide mediated endothelial control of vascular tone, and identifies the critical role of endothelial mitochondrial energy production in fueling perfused blood vessel function.

## 1. Introduction

Endothelial cells are one of the most abundant mammalian, non-blood cells in the body [1,2], and they form the inner lining of all blood vessels. In most adult tissue, they exist in a ‘quiescent’ state characterised by minimal or absent migration and proliferation. Indeed, human coronary endothelial cells *in vivo* have a slow replication rate, with renewal of the entire population taking approximately six years [3]. Whilst endothelial cell replication in mature blood vessels is negligible, the cells are highly active. Although paradoxically called ‘quiescent’, these endothelial cells play a crucial role in controlling blood flow by regulating blood vessel contraction and dilation. They also regulate the movement of fluid, metabolites, and cells between the bloodstream and body tissue. Impaired endothelial cell function precipitates, aggravates, and reinforces cardiovascular diseases such as atherosclerosis and hypertension. However, little attention has been paid to how normal endothelial cell function is regulated and this has limited our understanding of the progression of cardiovascular disease.

Alongside negligible replication rates, endothelial cells in perfused blood vessels are also considered to have a low basal metabolism [4]. Nevertheless, endothelial function requires energy. In most cells, the bulk of energy is provided by mitochondria in the form of ATP. But under oxygen limiting conditions, cells are forced to rely on glycolysis alone for ATP production. Remarkably, despite endothelial cells having abundant access to high concentrations of oxygen in the blood, four decades of reports have indicated that they use glycolysis for ATP production rather than mitochondrial oxidative phosphorylation [5–8]. The preference for glycolysis is proposed to allow ATP to be generated: 1) at a faster rate; 2) in the absence of oxygen; 3) to facilitate oxygen transfer to extravascular cells; and 4) to protect endothelial cells from reactive oxygen species. Together with observations that mitochondria occupy a smaller fraction of the volume of endothelial cells than more energetic hepatocytes/cardiac myocytes (~5% vs ~30%) [9,10], these proposals are used to justify the assumption that mitochondrial respiration is dispensable for endothelial physiology [11–15].

Nevertheless, work in the field of endothelial cell metabolism has intensified in recent years. This is particularly true for the study of angiogenesis, where a number of important studies have highlighted how endothelial cell glycolysis drives vascular function (see, for example, among a deluge of review articles those by Falkenberg *et al*. (2019) and Li *et al*. (2019) [16,17]). This evidence of a relationship between metabolism and function has led to a reappraisal of mitochondrial energy production in endothelial cells [18]. For example, global pharmacological inhibition and conditional, endothelial-specific knockout of the mitochondrial respiratory chain complex III each inhibit endothelial-driven wound healing [19,20]. Other studies demonstrate that mitochondrial respiration regulates fatty acid transport across the vascular wall [21], and is necessary for angiogenesis [22–25]. Together, these studies suggest that endothelial mitochondria play an important role in vascular function. However, the significance of mitochondrial metabolism has rarely been studied outside of the context of proliferative endothelial cells in angiogenesis or pulmonary hypertension [26], in which the contribution of the organelle may be diminished. Moreover, almost no attention has been given to the energetic requirements of perfused blood vessel endothelial cells.

In the present study, we tested the necessity of mitochondrial ATP production for the obligatory role of endothelial cells in the relaxation of arterial smooth muscle [27]. We found that preventing mitochondrial production of ATP at complex V increases blood vessel contraction and inhibits endothelium-dependent blood vessel relaxation. The requirement for mitochondrial ATP production appears to be independent of sex, species, or vascular bed. Our results provide evidence that mitochondrial energy production in endothelial cells critically regulates blood vessel diameter. Moreover, mitochondrial energy production is far more intimately linked to endothelial cell physiology than is currently appreciated.

## Results

### Mitochondrial ATP fuels nitric oxide-mediated endothelial control of vascular tone

To investigate the role of mitochondrial energy metabolism in blood flow control, we used a single-photon microscope optimized to visualize endothelial cells and detect increases or decreases in vascular tone in small mesenteric arteries (Figure 1A-B) [28,29]. Vessel tone was monitored as arteries were stimulated with the α-adrenergic receptor agonist, phenylephrine, and then the muscarinic receptor agonist, acetylcholine (ACh), in the absence and presence of the mitochondrial ATP synthase (complex V) inhibitor, oligomycin (2.4 μM; Figure 1C-D, Video 1). The concentration of phenylephrine (~2 μM) was titrated to achieve a ~20% increase in artery tone in control conditions. At the plateau phase of the constriction, ACh (10 μM) evoked a robust dilation. Following washout, vasomotor responses were reproducible. Significantly, subsequent inhibition of mitochondrial ATP production using oligomycin reversed relaxations to ACh. The reversal of ACh-evoked dilation by oligomycin was observed in arteries from both male and female rats (Figure 1E-F), and neither global (TEMPO, 100 μM) nor mitochondrially-targeted (mitoTEMPOL, 50 μM) scavenging of reactive oxygen species prevented the reversal of ACh-evoked vasodilation by oligomycin (Figure 2A-D). Additionally, oligomycin did not prevent endothelium-independent vasodilation to sodium nitroprusside (SNP, 100 μM) whether in the absence (Figure 1C-E) or presence of the reactive oxygen species scavengers (Figure 2A-D). The functional impairment induced by mitochondrial complex V inhibition was mimicked by inhibiting respiration at complex I using rotenone (500 nM; Figure 2E-F). These data demonstrate that mitochondrial ATP regulates endothelium-dependent control of vascular tone.

**Figure 1 –.**
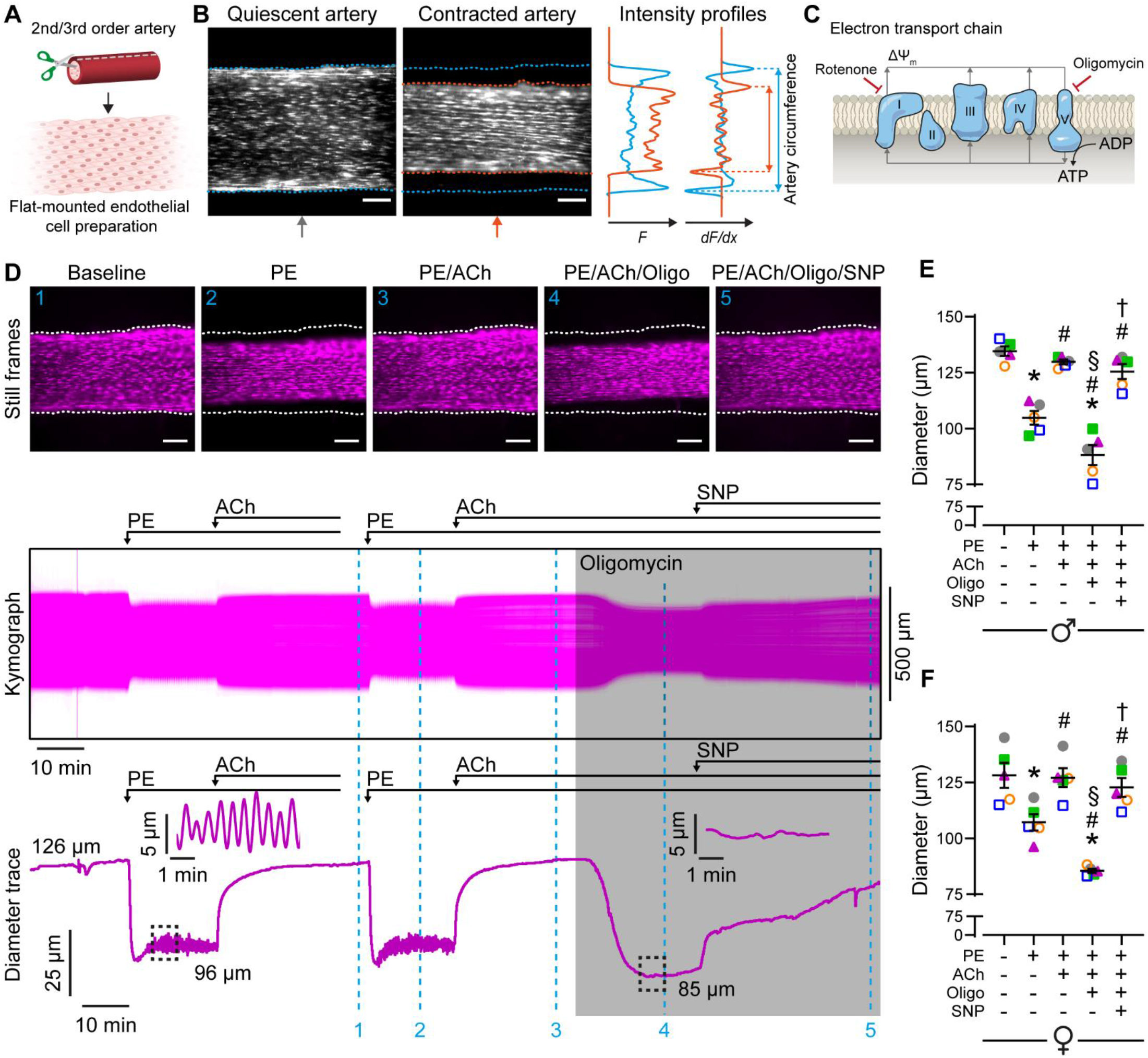
Inhibiting oxidative phosphorylation reverses endothelium-dependent vasodilation. (A) Schematic of dissection procedure for *en face* endothelial cell imaging of flat-mounted blood vessels. Blood vessels are cut open and mounted in a low-volume imaging chamber with the endothelium facing up. Agents that alter vascular activity are delivered using a continuous perfusion system. (B) Still frame images of mesenteric artery endothelial cells before and during stimulation of smooth muscle cell contraction. The position of the artery edges is determined using the derivative of intensity profiles that bisect the artery. C) Scheme depicting the sites of action for mitochondrial inhibitors used in the study. (D) Still frame images, kymograph, and calculated diameter trace showing the effect of oligomycin (2.4 μM) on repeatable vasodilation to acetylcholine (ACh, 10 μM) in arteries contracted with phenylephrine (PE, concentration adjusted to achieve ~20% contraction). Sodium nitroprusside (SNP; 100 μM) was used to test endothelium-independent vasodilation. Data shown in Video 1. (E-F) Summary data (mean ± SEM overlaid) showing the effect of oligomycin on vasomotor responses in mesenteric arteries from male (E) and female (F) rats. Each color-coded set of data points represents repeat measurement from a single artery (n = 5, each from a different animal). The dataset in D is from a female rat and is shown in F as magenta triangles. Significance markers indicates statistical significance (p < 0.05) using repeated measures one-way ANOVA with Tukey’s test for multiple comparisons (* vs baseline; # vs PE; § vs PE/ACh; † vs PE/ACh/oligomycin). Image scale bars = 100 μm.

**Figure 2.**
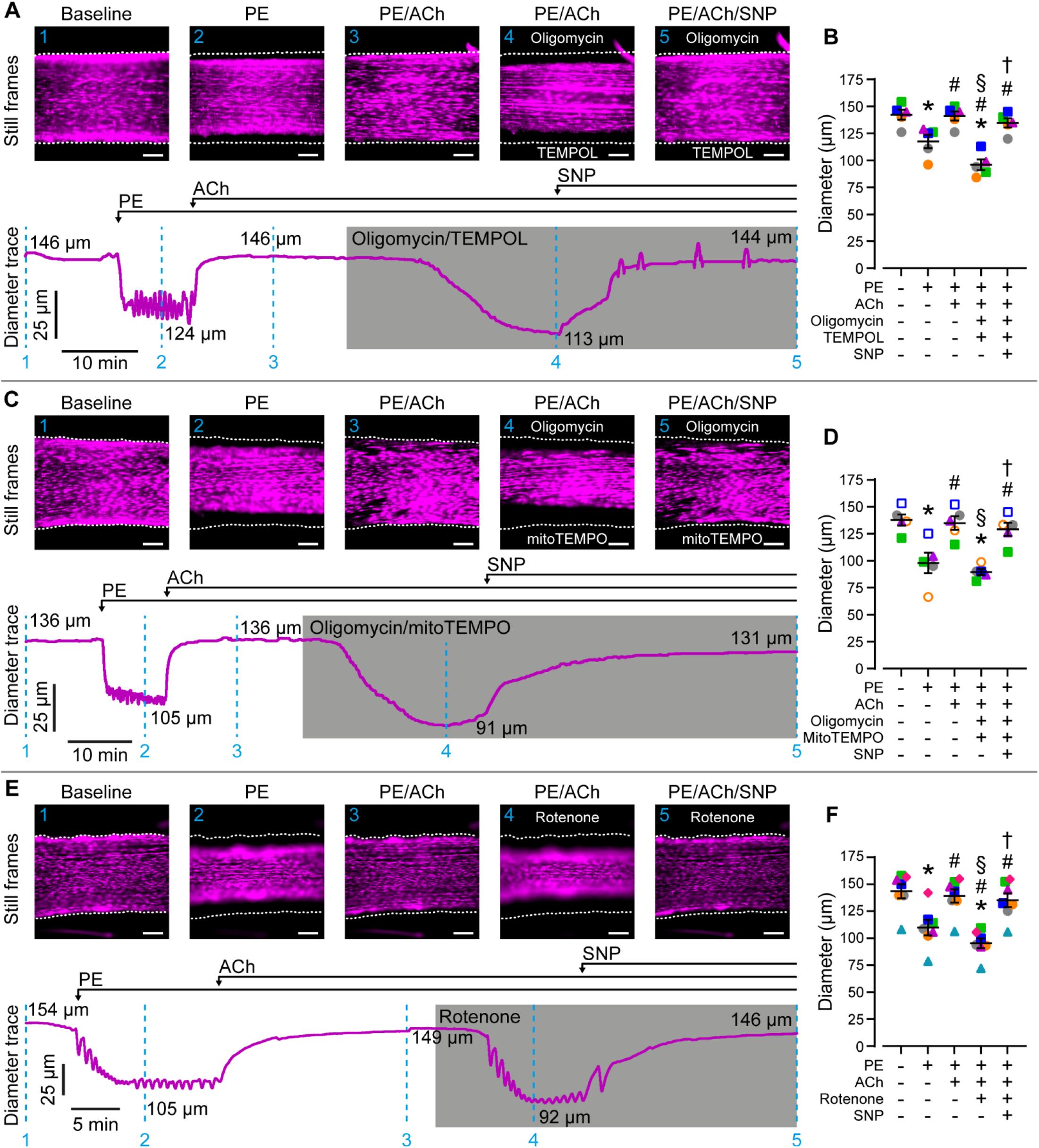
Scavenging reactive oxygen species does not rescue endothelium-dependent vasomotor control following mitochondrial oxidative phosphorylation impairment. (A-F) Still frame images, diameter traces, and summary data showing:1) the effect of global (A-B) and mitochondrial-targeted (C-D) reactive oxygen species scavenging on the reversal of acetylcholine-evoked (ACh, 10 μM) vasodilation following mitochondrial ATP synthase impairment (using oligomycin; 2.4 μM); and 2) the effect of mitochondrial complex I inhibition (using rotenone; rote, 500 nM) on ACh-evoked vasodilation (E-F). Arteries contracted with phenylephrine (PE, concentration adjusted to achieve ~20% contraction). Summary data (mean ± SEM overlaid) are colored to highlight measurements from a single artery (n = 5-7, each from a different animal). Data in A,C,E are each shown in corresponding summary data as magenta triangles. Significance markers indicate p < 0.05 using repeated measures one-way ANOVA with Tukey’s test for multiple comparisons (* vs baseline; # vs PE; § vs PE/ACh; † vs PE/ACh/oligomycin). Image scale bars = 100 μm.

To examine whether the mitochondrial inhibitors prevented endothelial cell vasodilator signaling or induced smooth muscle cell contraction or both, we preincubated arteries with oligomycin and reexamined vascular reactivity. When applied under resting conditions (in the absence of vasoactive stimuli) oligomycin did not evoke vascular contraction (Figure 3A). However, subsequently, phenylephrine-induced artery constriction was enhanced and ACh-evoked dilation was impaired (Figure 3A-D). Again, oligomycin did not prevent endothelium-independent relaxations to SNP (Figure 3D). The effects of oligomycin (increased vessel constriction, decreased dilation) mimic those induced by mechanical ablation of the endothelial cell layer [28]. These data indicate that endothelial cells are the site of action of the mitochondrial inhibitors, and suggest that mitochondrial generation of ATP underlies the ability of endothelial cells to: 1) oppose vascular constriction; and 2) to promote ACh-evoked vascular relaxation.

**Figure 3.**
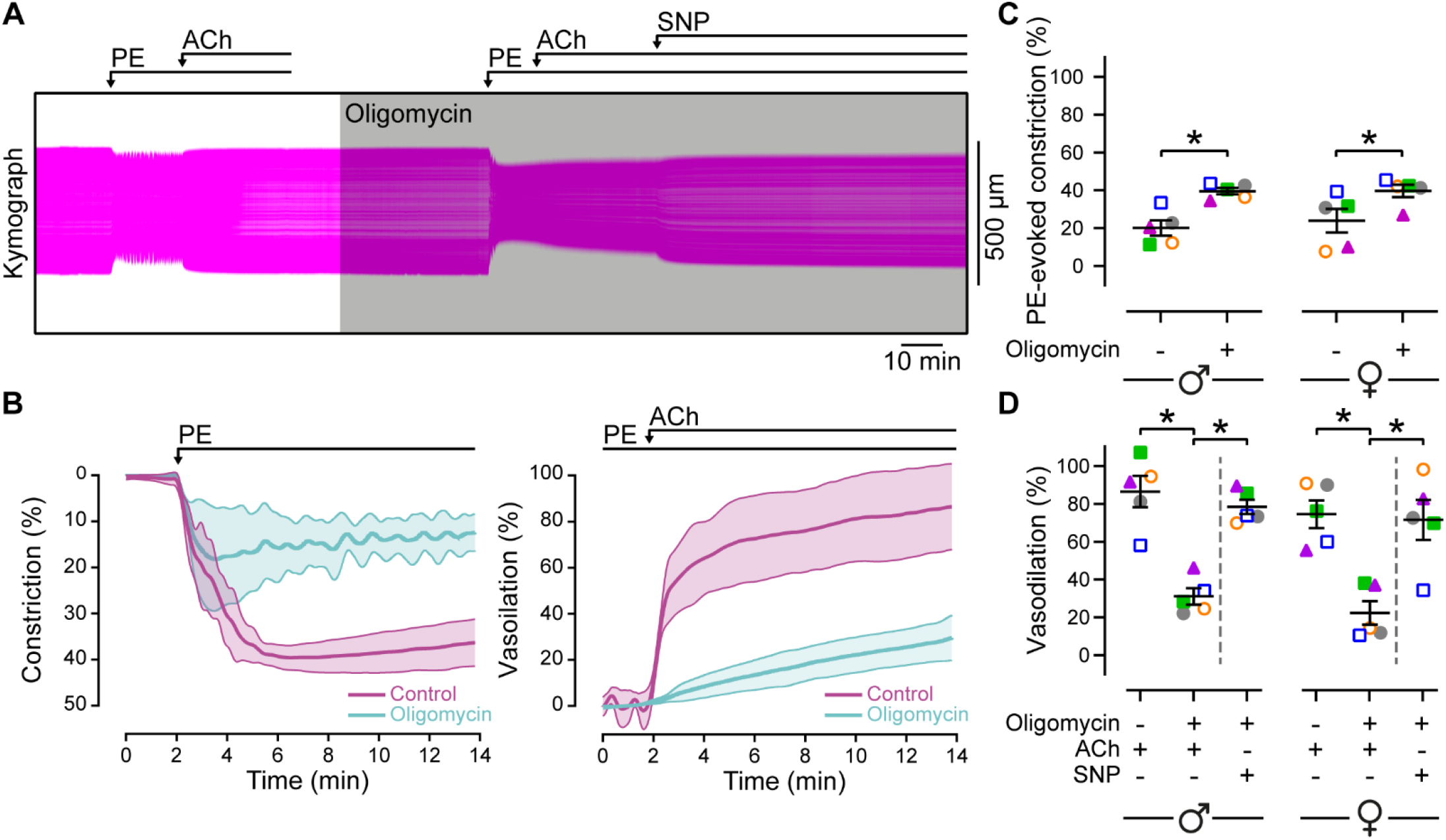
Oxidative phosphorylation is required for endothelial modulation of vascular tone. Representative kymograph showing the effect of oligomycin (2.4 μM) preincubation on phenylephrine (PE)-induced constriction and acetylcholine (ACh)-evoked (10 μM) dilation in mesenteric arteries (PE). Constriction is expressed as % of resting diameter. Sodium nitroprusside (SNP; 100 μM) was used to test endothelium-independent vasodilation. (B) Mean ± SEM time courses of PE-evoked constriction (left) and ACh-evoked dilation (right) before and after incubation with oligomycin (n = 5). Constriction data expressed as % of initial diameter, vasodilation data expressed as percentage of maximal relaxation (constricted diameter to resting diameter). (C-D). Paired summary data plots (mean ± SEM overlaid) showing the effect of oligomycin on artery constriction (C) and vasodilation (D) in male and female rats. PE concentration titrated to achieve ~20% constriction in the absence of oligomycin. Individual data points are color-coded to indicate measurements from a single artery (each from a different animal). The dataset in A is from a male rat and is shown in C & D as grey circles. * indicates statistical significance (p < 0.05) using paired t-test (C) or repeated measures one-way ANOVA with Dunnett’s test for multiple comparisons (D).

Endothelium-dependent opposition to vascular tone and relaxation to ACh occurs mainly via the production of nitric oxide [28,30]. Endothelial cells may also evoke nitric oxide-independent vascular relaxation via the activation of intermediate (IK) and small (SK) conductance Ca^2+^-activated K^+^ channels, which mediate smooth muscle cell hyperpolarization. As TRPV4 channels are known to modulate endothelial control of vascular tone via this endothelial-derived hyperpolarization pathway [31,32], we also pharmacologically activated TRPV4 channels after exposure to oligomycin (Figure 4A-B). Whilst, inhibition of mitochondrial ATP production reversed ACh-evoked vasodilation, it did not prevent concentration-dependent relaxations to the specific TRPV4 channel agonist, GSK1016790A. These data suggest that mitochondrial ATP modulates nitric oxide-mediated control of vascular tone, but not vasodilation arising from endothelial-derived hyperpolarization pathways.

**Figure 4.**
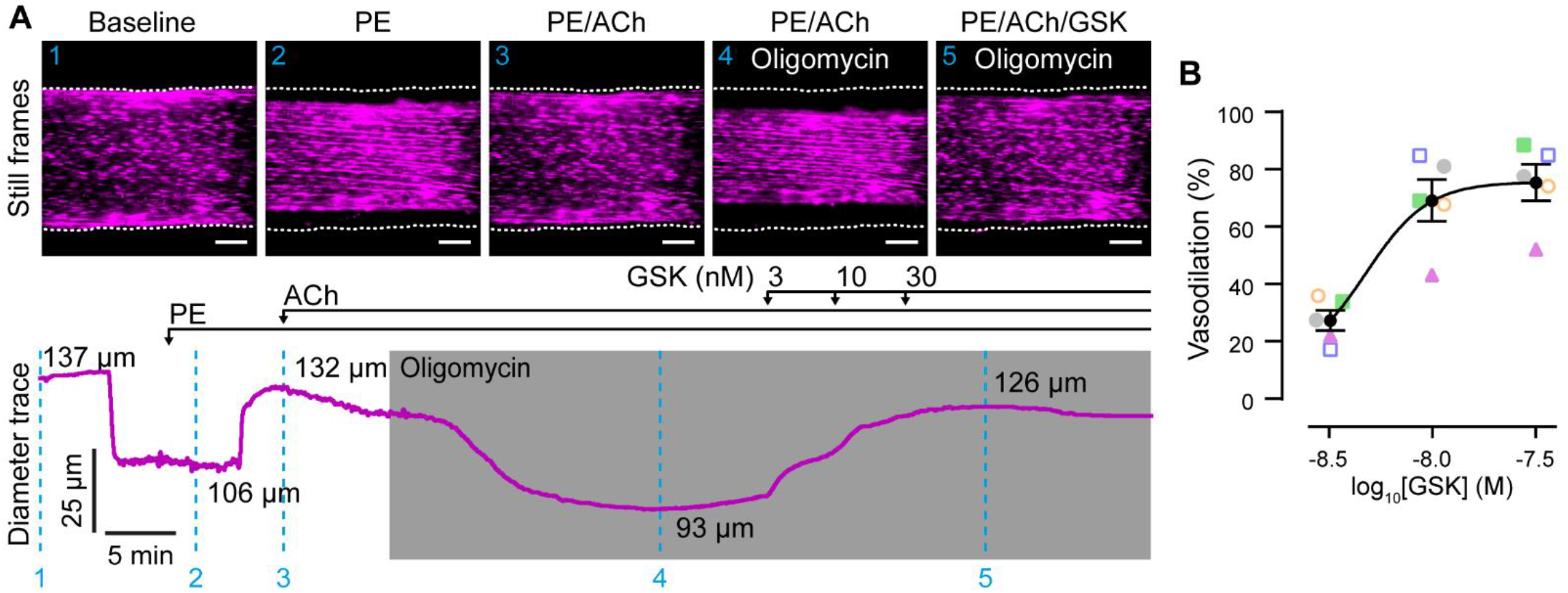
TRPV4-evoked vasodilation is preserved after inhibition of the mitochondrial ATP synthase. (A) Representative still frame images and diameter trace showing the concentration dependence of TRPV4-mediated vasodilation following pharmacological inhibition of the mitochondrial ATP synthase using oligomycin (2.4 μM). Arteries were first constricted with phenylephrine (PE, concentration adjusted to achieve ~20% contraction) and then activated using acetylcholine (ACh, 10 μM). ACh-evoked vasodilation was reversed, using oligomycin, and arteries were then challenged with increasing concentrations of the specific TRPV4 channel agonist, GSK1016790A (GSK). (B) Summary data showing the concentration dependence of TRPV4-mediated vasodilation in arteries with pharmacologically impaired mitochondrial ATPase function. Vasodilation data expressed as percentage of maximal relaxation (constricted diameter to resting diameter). Each colored data point represents measurements from a single artery (each from a different animal, n = 5). The dataset in A is shown in B as magenta triangles. Summary data are mean ± SEM; * indicates statistical significance (p < 0.05, versus PE/ACh/oligomycin) using repeated measures one-way ANOVA with Dunnett’s test for multiple comparisons. Image scale bars = 100 μm.

### 2.2. Mitochondrially derived ATP facilitates IP_3_-mediated endothelial cell calcium signalling

We next asked, how does mitochondrial ATP facilitate nitric oxide-mediated vasodilation? The Ach-nitric oxide vasodilator pathway is mediated via muscarinic (M3 subtype [33]), Gαq-dependent stimulation of phospholipase C and the subsequent hydrolysis of phosphatidylinositol-4,5-bisphosphate (PIP2) to produce IP_3_, Once liberated, IP_3_ activates IP_3_ receptors evoke, in turn, Ca^2+^ release from internal stores, Ca^2+^ influx via store-operated Ca^2+^ entry, and Ca^2+^-dependent activation of endothelial nitric oxide synthase [28,30]. Because functional mitochondria are required for basal (unstimulated) endothelial cell Ca^2+^ signaling [34], we speculated that mitochondrial ATP synthase control of nitric-oxide mediated vasodilator signaling (Figures 1–3) arises via the nucleotide’s control of IP_3_-mediated Ca^2+^ signaling.

To test this hypothesis, we used high resolution, wide-field single photon imaging to probe Ca^2+^ signaling in large populations of endothelial cells in intact arteries. Using a paired (before/after) experimental design, we examined the effect of mitochondrial ATP synthase impairment on basal (unstimulated), flow-evoked, and ACh-evoked endothelial cell Ca^2+^ activity, each of which are dependent on IP_3_-mediated Ca^2+^ release [29,30,34,35]. Oligomycin inhibited each of these three endothelial Ca^2+^ signaling modalities (Figures 5A-E, Video 2, Video 3, Video 4). Oligomycin also inhibited ACh-evoked Ca^2+^ signaling in freshly isolated endothelial cell sheets, which lack contacts with smooth muscle cells (Figure 6A-D, Video 5). Together with our observation that oligomycin did not inhibit intact artery contraction (Figure 3), these results demonstrate a direct effect of oligomycin on IP_3_-mediated endothelial cell Ca^2+^ signaling. In contrast, Ca^2+^ responses evoked by the TRPV4 Ca^2+^ influx channel agonist, GSK1016790A (20 nM), persisted in the presence of oligomycin (Figure 5D-E), indicating that TRPV4-mediated Ca^2+^ influx is unaffected by mitochondrial ATP synthase inhibition. Collectively, these data indicate that mitochondrial ATP facilitates IP_3_-mediated endothelial cell Ca^2+^ activity.

**Figure 5.**
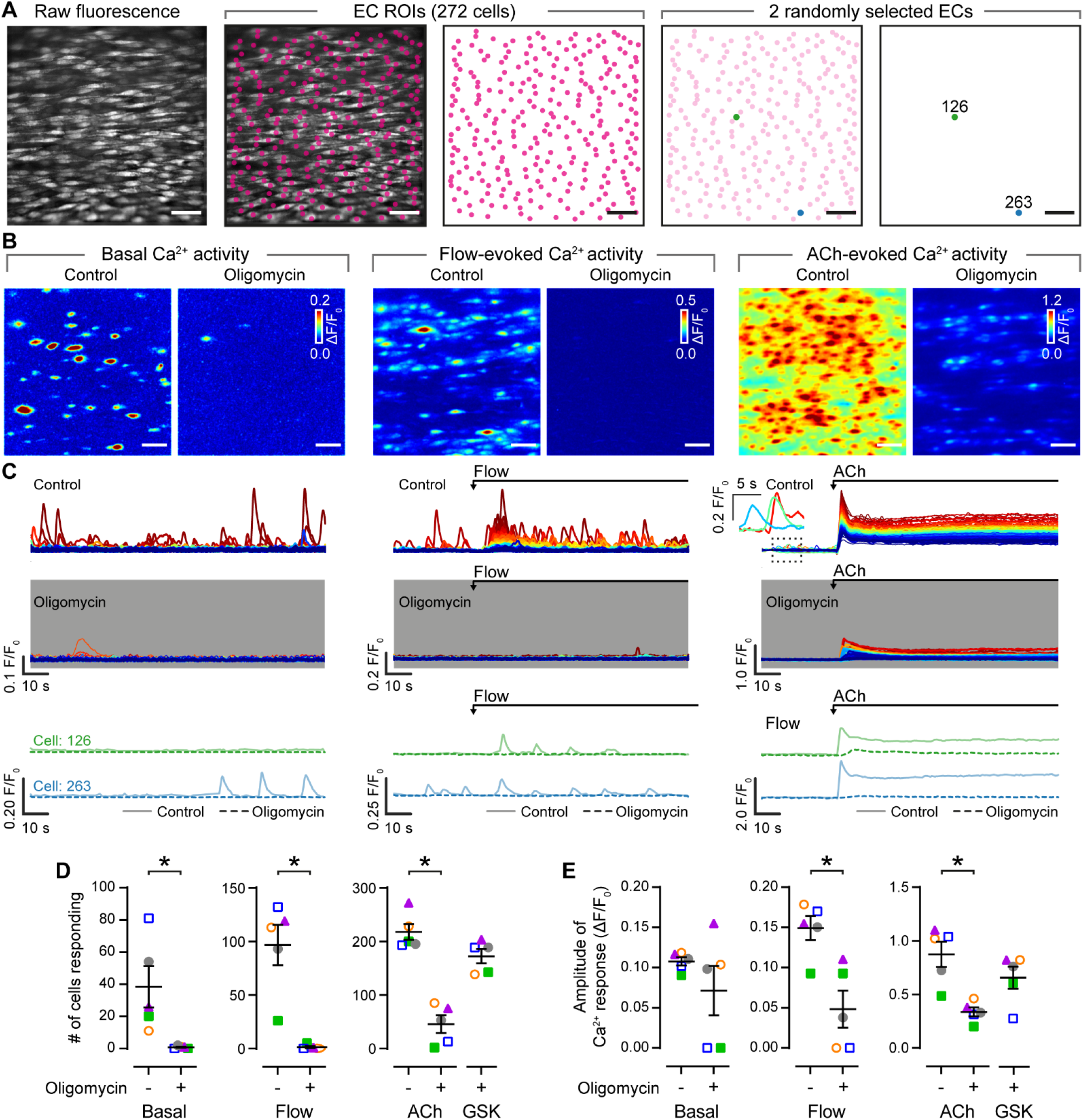
Endothelial calcium signaling requires functional mitochondrial ATP synthase. (A) Representative images of population level endothelial Ca^2+^ imaging. Regions of interest (ROIs) were automatically generated for all endothelial cells (ECs) visualized using Cal-520/AM (5 μM). (B-C) Ca^2+^ activity images (B) and corresponding single-cell Ca^2+^ signals (C) illustrating basal, flow-evoked, and acetylcholine (ACh)-evoked Ca^2+^ activity before (control) and after incubation with the mitochondrial ATPase inhibitor, oligomycin (2.4 μM). In B, Ca^2+^ images are pseudo colored maximum intensity projections of ΔF/F_0_ datasets (2-minute recordings). Images are grouped by stimulus and the same color scale is used for both images in each set. The same y-axis scale is used for Ca^2+^ traces within each set. In C, overlaid Ca^2+^ traces are colored by the magnitude of the response to ACh. All Ca^2+^ and traces show data from the field of endothelial cells in A. D-E) Summary data showing the effect of oligomycin on basal, flow-evoked, and ACh-evoked (right) endothelial Ca^2+^ activity. At the end of each experiment, Ca^2+^ responses to the TRPV4 antagonist, GSK1016790A (GSK; 20 nM) were recorded. Each color-coded set of data points represents repeat measurements from a single artery (n = 5, each from a different animal). The dataset in A-C is summarized in D-E as magenta triangles. The blue outlined square datapoints are shown in Videos 2-4. Summary data are mean ± SEM; * indicates statistical significance (p < 0.05) using paired t-test (basal/flow) or repeated measures one-way ANOVA with Dunnett’s test (agonist-evoked activity, vs Ach in absence of oligomycin). Scale bars = 50 μm.

**Figure 6.**
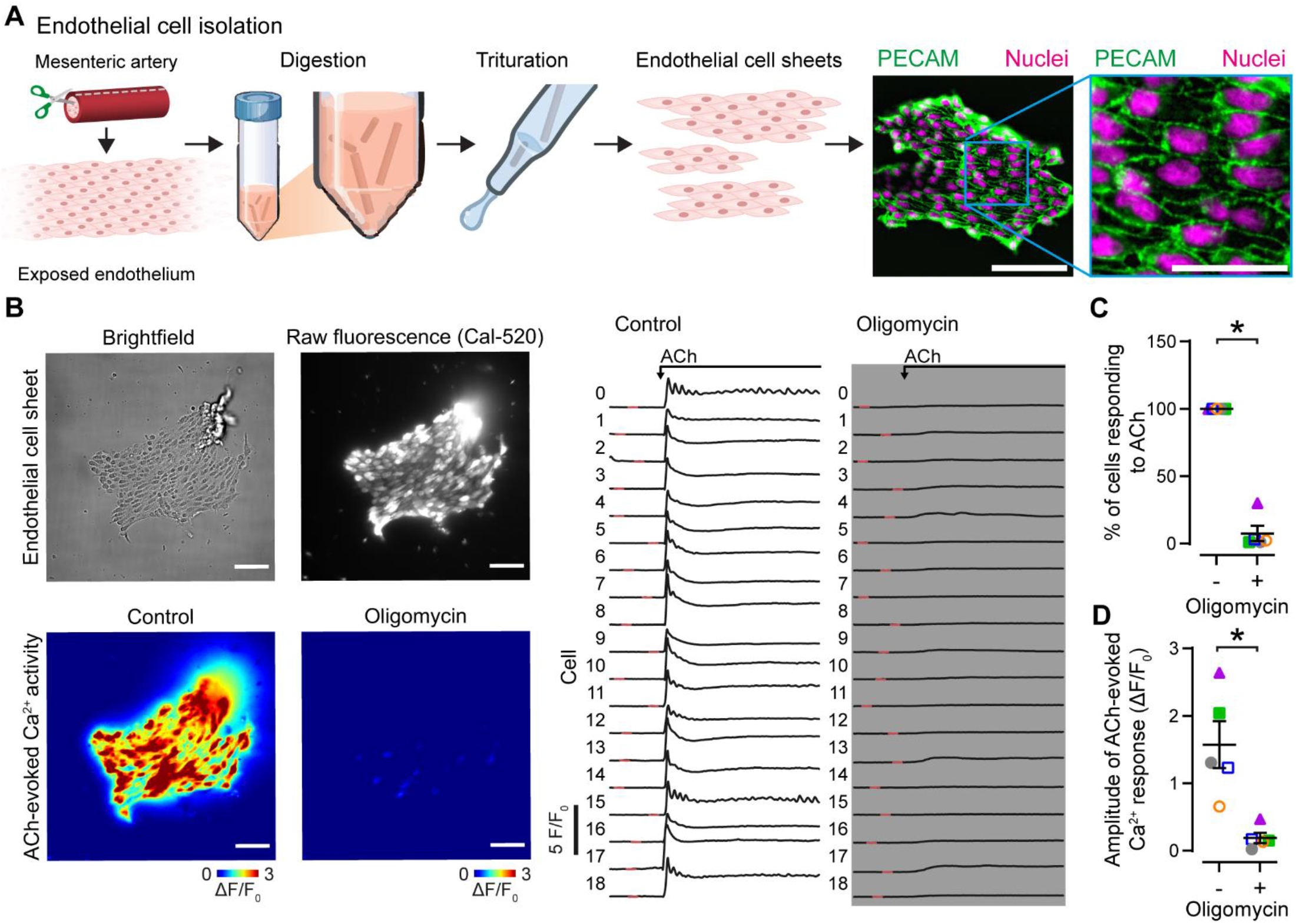
Mitochondrial ATP is essential for calcium signaling in isolated endothelial cells. (A) Schematic of endothelial cell isolation procedure and fluorescence image of an isolated endothelial cell sheet stained with the nuclear stain, DAPI, and for the endothelial specific marker, platelet endothelial cell adhesion molecule (PECAM). Scale bar = 50 μm (25 μm, inset). (B) Representative brightfield, fluorescence and Ca^2+^ activity images, and corresponding single cell traces illustrating acetylcholine (ACh)-evoked Ca^2+^ activity in isolated endothelial cell sheets before and after inhibition of the mitochondrial ATP synthase using oligomycin (2.4 μM). Ca^2+^ images are pseudo colored maximum intensity projections of ΔF/F_0_ datasets (2-minute recordings). (C-D) Summary data showing the effect of oligomycin on ACh-evoked Ca^2+^ activity in isolated endothelial cell sheets. Each color-coded set of data points represents repeat measurements from a single artery (n = 5, each from a different animal). The dataset in A-C is summarized in D as magenta triangles and shown in Video 5. Summary data are mean ± SEM; * indicates statistical significance (p < 0.05) using paired t-test. For presentation purposes, the data in C is shown as the percentage of cells responding to ACh, statistical tests were performed on the original cell counts. Scale bars = 50 μm (25 μm, inset in A).

To further investigate mitochondrial control of IP_3_-mediated Ca^2+^ signaling, we dual-loaded intact artery endothelial cells with a caged form of IP_3_ and the Ca^2+^ indicator, Cal-520/AM. This strategy allowed us to activate IP_3_ receptors by photo-releasing IP_3_ using an ultraviolet laser, bypassing phospholipase C activation and IP_3_ production, whilst performing simultaneous single-photon Ca^2+^ imaging (Figure 7A-B). This allowed us to distinguish between mitochondrial control of IP_3_ production and IP_3_ receptor activation. Photo-stimulated Ca^2+^ activity was inhibited following mitochondrial ATP synthase impairment (Figure 7C-D, Video 6). Significantly, oligomycin does not alter the Ca^2+^ store content in native endothelial cells [34]. Thus, mitochondrial ATP is required for the activation of IP_3_ receptors.

**Figure 7.**
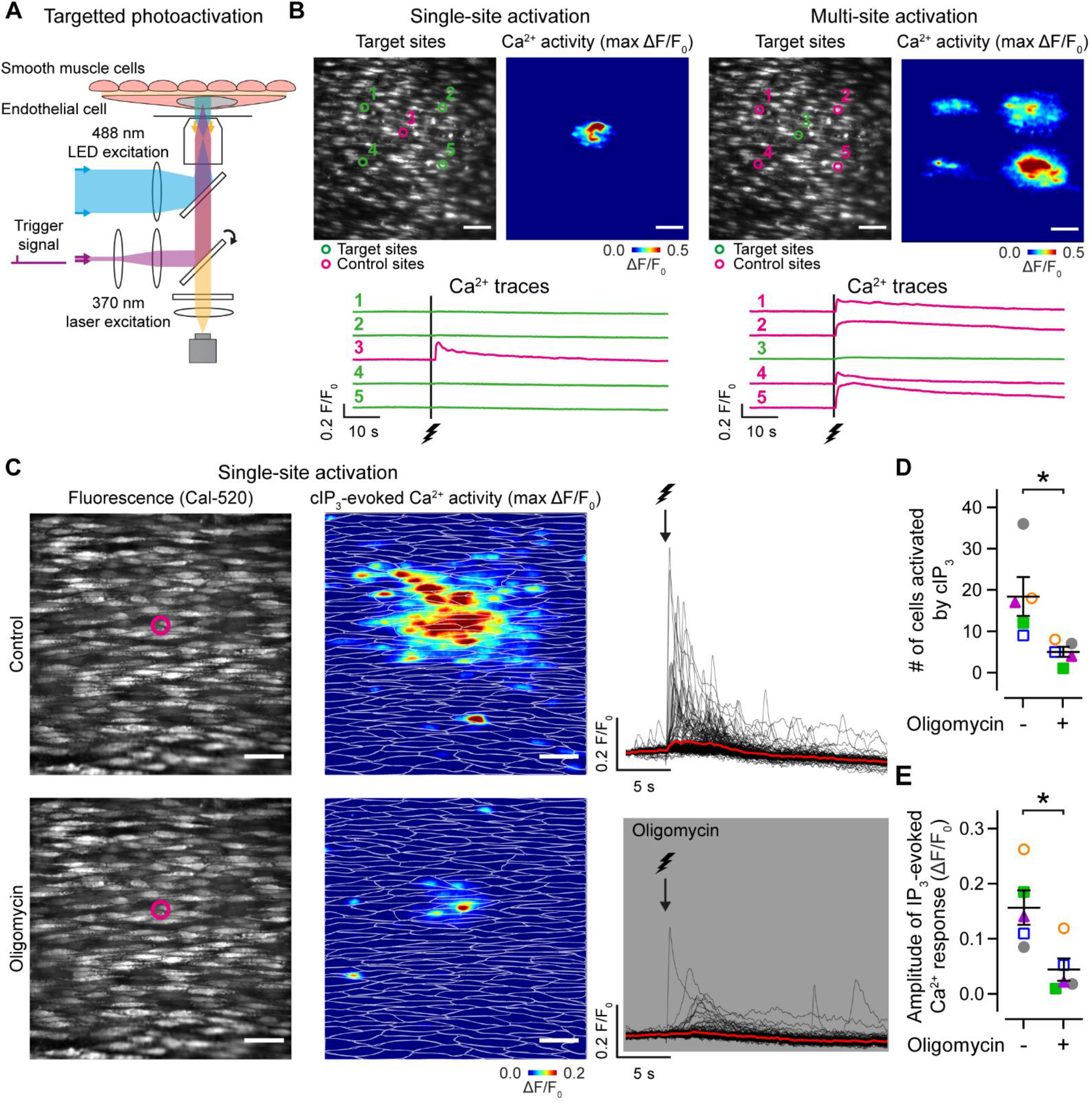
Direct activation of endothelial IP_3_ receptors requires mitochondrial ATP. (A-B) Schematic of singlephoton microscopy with targeted endothelial cell photoactivation. (B) Example fluorescence and Ca^2+^ activity images, and corresponding Ca^2+^ traces illustrating single- and multi-site activation of endothelial cell IP_3_ receptors via photolysis of caged IP_3_ (cIP_3_). Endothelial cells were dual loaded with the Ca^2+^ indicator, Cal-520 (10 μM), and cIP_3_. Green circles denote photoactivation target regions, magenta circles indicate control (no stimulation) regions. Photolysis of caged IP_3_ evoked robust Ca^2+^ responses. (C) Representative fluorescence and Ca^2+^ activity images, and corresponding single-cell Ca^2+^ traces illustrating single-target Ca^2+^ responses before (top, control) and after (bottom) inhibition of the mitochondrial ATP synthase with oligomycin (2.4 μM). (D-E) Summary data showing the effect of oligomycin on IP_3_-evoked endothelial Ca^2+^ activity. Each color-coded set of data points represents repeat measurements from a single artery (n = 5, each from a different animal). The dataset in C is summarized in D & E as magenta triangles. See Video 6. Summary data are mean ± SEM; * indicates statistical significance (p < 0.05) using paired t-test. All image scale bars = 50 μm.

### 2.3. Mitochondrial oxidation of pyruvate drives mitochondrial control of endothelial function

Since mitochondrial ATP is required for endothelial cell IP_3_-mediated Ca^2+^ signaling, we reasoned that inhibiting the transport of pyruvate - the major fuel for mitochondrial energy production – would also impair the endothelial Ca^2+^ response to ACh. Mitochondrial pyruvate metabolism requires that the substrate (pyruvate) is transported into the mitochondrial matrix by the mitochondrial pyruvate carrier (MPC) [36]. Inhibition of mitochondrial uptake of pyruvate using either UK5099 (50 μM) or MSDC-0160 (mitoglitazone; 10 μM) decreased ACh-evoked endothelial cell Ca^2+^ signaling (Figure 8A-C). The action of these inhibitors demonstrate that pyruvate oxidation is an important energy pathway in ‘quiescent’ endothelial cells of the fully formed vascular wall.

**Figure 8.**
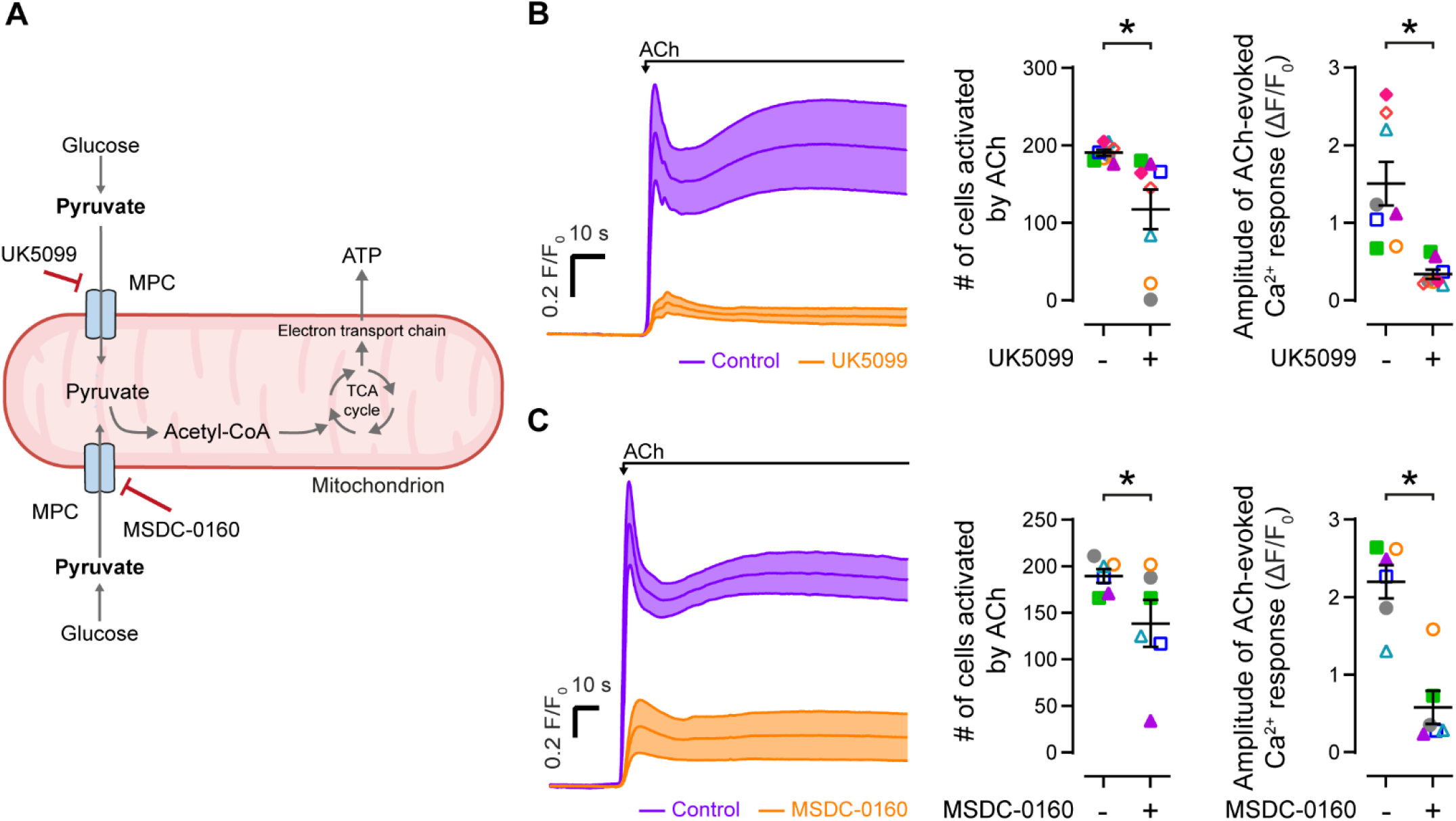
Limiting mitochondrial pyruvate transport inhibits endothelial cell calcium signaling. (A) Scheme depicting the sites of action for mitochondrial pyruvate transport inhibitors. (B-C) Mean ± SEM time courses (before and after), and summary data plots showing the effect of the mitochondrial pyruvate carrier inhibitors, UK5099 (B; 50 μM; n = 8) and MSDC-0160 (mitoglitazone; 10 μM; n = 6) on acetylcholine (ACh)-evoked (10 μM) endothelial Ca^2+^ activity. Each color-coded set of data points represents repeat measurements from a single artery (each from a different animal). Summary data are mean ± SEM; * indicates statistical significance (p < 0.05) using paired t-test.

### 2.4. Mitochondrial ATP is a pan-endothelial regulator of vascular function

As endothelial cells exhibit a bewildering degree of heterogeneity among and within vascular beds [37,38], we asked one final question: is the control of nitric oxide vasodilator signaling by mitochondrial ATP a universal feature of endothelial cell function? To answer this question, we examined muscarinic receptor-mediated endothelial Ca^2+^ activity in cerebral, coronary, and renal microvessels of the rat (Figure 9A-B). We also examined endothelial Ca^2+^ signaling in mouse mesenteric arteries (Figure 9C-D). Regardless of vascular bed, sex, or species, IP_3_-mediated Ca^2+^ signaling required functional mitochondrial ATP synthase (Figure 9). Thus, mitochondrial ATP production is a ubiquitous energy pathway in quiescent endothelial cells of mature blood vessels.

**Figure 9.**
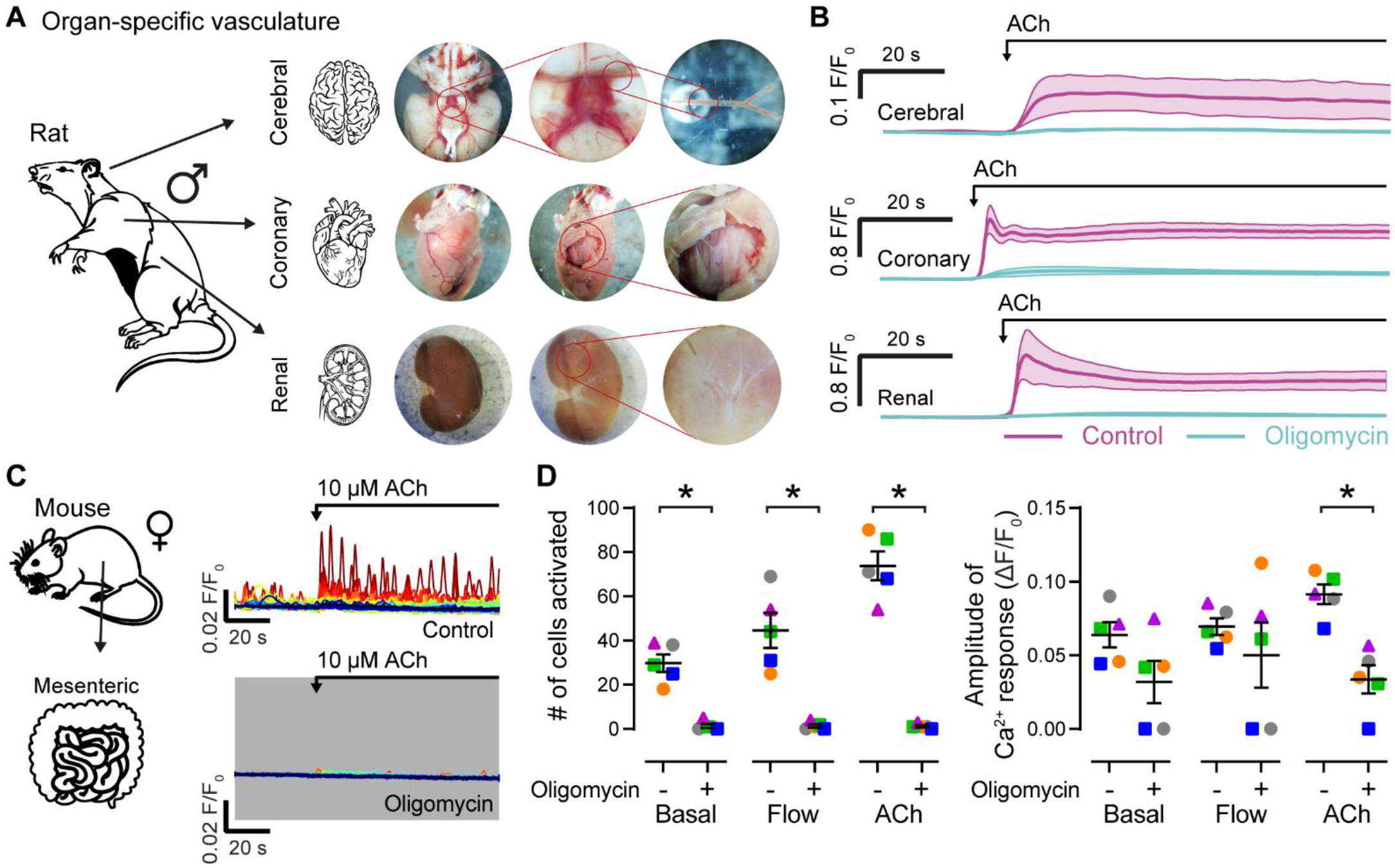
Mitochondrial ATP is a pan-endothelial regulator of calcium signaling. (A-B) Isolation procedure (A) and mean ± SEM Ca^2+^ signals (B; before and after) showing the effect of the mitochondrial ATPase inhibitor, oligomycin (2.4 μM), on acetylcholine (ACh)-evoked (10 μM) endothelial Ca^2+^ activity in rat cerebral (n = 3), coronary (n = 3), and renal (n = 3) microvessels. (C-D) Example single-cell Ca^2+^ traces (C; before and after), and paired summary data plots (D) showing the effect of oligomycin (2.4 μM) on basal, flow-evoked, and ACh-evoked (10 μM) endothelial cell Ca^2+^ signaling in mouse mesenteric arteries. Each color-coded set of data points represents repeat measurements from a single artery (n = 5, each from a different animal). The dataset in C is summarized in D as grey circles. Summary data are mean ± SEM (n = 5); * indicates statistical significance (p < 0.05) using paired t-test.

## 3. Discussion

### 3.1. Arterial endothelial cells use oxidative phosphorylation for blood flow control

Mammalian cells generally rely on mitochondrial respiration to fuel bio-energetic processes, but endothelial cells are considered to be an exception to this rule. Rather than relying on mitochondrial ATP production, endothelial cells are thought to meet approximately 85% of their energy demand via glycolysis [6–8,39]. These estimations have emerged from studies examining the angiogenic potential of cultured proliferative/migratory endothelial cells as a model of angiogenesis. However, the majority of endothelial cells in perfused blood vessels are neither proliferative nor migratory (e.g., less than 2% of endothelial cells in liver and spleen have a proliferative phenotype [40]). Nevertheless, it is assumed that so-called ‘quiescent’ endothelial cells are glycolytic and that mitochondrial-derived energy plays little role in regulating endothelial function. In stark contrast to this assumption, we provide clear evidence that mitochondrial ATP production is crucial for the most widely-known function of endothelial cells in mature blood vessels - the control of artery diameter.

Endothelial cell control of artery diameter is exemplified by the vascular response (vasodilation) to the classical neurotransmitter, ACh [27], which is released by endothelial cells in response to fluid flow [35,41]. When activated by ACh, endothelial muscarinic receptors stimulate the production of nitric oxide by endothelial nitric oxide synthase. Once produced, nitric oxide diffuses to adjacent smooth muscle cells to cause smooth muscle cell relaxation and vasodilation. In parallel, ACh may also initiate the spread of a hyperpolarizing electrical signal (endothelium-dependent hyperpolarization), that also acts to relax adjacent smooth muscle cells. When mitochondrial ATP production was prevented, a rapid and complete loss of ACh-evoked endothelium-dependent vasomotor control occurred – endothelial cells failed to oppose vascular tone, and they failed to initiate vasodilation. As rat mesenteric artery vasodilation is predominantly mediated by nitric oxide signaling [28–30], these defects are probably not due to a failure in the endothelium-derived hyperpolarization pathway. Our finding that the TRPV4 channel agonist and activator of endothelium-derived hyperpolarization [35,41], GSK1016790A, elicited robust, concentration-dependent vasodilation following mitochondrial impairment adds support to this conclusion. Furthermore, as smooth muscle cell contraction and endothelium-independent vasodilation were not impaired following the inhibition of mitochondrial respiration, the defect in nitric oxide signaling likely arises due to a direct effect on endothelial cells and not on smooth muscle cells. Our conclusions are consistent with observations that endothelial mitochondria do generate ATP [5–8,42–47] and that the organelle increases ATP production when challenged [48–50],

### 3.2. Mitochondrial ATP is required for endothelial cell calcium activity

The impairment of endothelial vasodilator signaling after mitochondrial ATP synthase inhibition appears to be due to the inability of the cells to sustain the nitric oxide signaling pathway. Endothelial nitric oxide synthase is a Ca^2+^-dependent enzyme that is activated by intracellular Ca^2+^ signals. In unstimulated endothelial cells, spontaneous Ca^2+^ release events via IP_3_ receptors give rise to a basal level of nitric oxide that opposes vascular tone [29]. Mechanical and chemical stimuli each amplify this basal Ca^2+^ signaling modality, whilst also eliciting Ca^2+^ entry, to stimulate nitric oxide production and promote vasodilation. We observed a severe disruption of basal, flow-, agonist-, and directly-evoked IP_3_-mediated Ca^2+^ signaling in respiration-deficient endothelial cells in intact artery endothelium and in freshly isolated sheets of endothelial cells. Preventing the transport of pyruvate by targeting the mitochondrial pyruvate carrier also inhibited endothelial cell Ca^2+^ signaling. Since ACh-evoked Ca^2+^ entry occurs via store-operated mechanisms, a consequence of IP_3_-mediated Ca^2+^ release, the dependence of endothelial vasomotor control on mitochondrial ATP appears to arise via the nucleotides control of endothelial cell Ca^2+^ release.

Intriguingly, and in line with our vascular tone experiments, mitochondrial ATP production was not required for Ca^2+^ influx via TRPV4 channels. This result provides further support for our conclusion that the effect of mitochondrial ATP synthase inhibition is due to modulation of Ca^2+^ release pathways, rather than a direct effect on Ca^2+^ entry. As oligomycin does not cause a reduction in endothelial cell Ca^2+^ store content [34], it is likely that this control arises via IP_3_ receptor activity. Two mechanisms may explain how mitochondrial ATP modulates IP_3_ receptor-mediated Ca^2+^ signaling to promote endothelial vasodilator signaling. First, the generation of IP_3_ and its precursors by phospholipase C and phosphoinositide kinases require millimolar concentrations of intracellular ATP [51–54]. Second, by binding specifically to an ATP-binding site on the IP_3_ receptor, micromolar concentrations of ATP may sensitize the channel to promote Ca^2+^ release [55–57]. Thus, in the absence of sufficient cytosolic ATP, IP_3_ receptor-mediated Ca^2+^ signaling may fail. Because of our finding that Ca^2+^ responses evoked by direct activation of IP_3_ receptors require a functional mitochondrial ATP synthase, we speculate that the mitochondrial ATP requirement resides at the IP_3_ receptor. Going forward, it will be important to determine if a loss of mitochondrial-derived ATP also prevents IP_3_ production.

As well as fueling the molecular machinery responsible for intracellular Ca^2+^ signals, mitochondria regulate Ca^2+^ release/influx through the generation of reactive oxygen species [58–60]. The organelle also buffers cytosolic Ca^2+^ directly by accumulating/releasing the ion. Neither of these mitochondrial activities are likely to explain the present findings. With regards to the former, reactive oxygen species are produced as a consequence of normal electron transport chain function and during mitochondrial dysfunction. For example, inhibition of mitochondrial complex V by the ATPase Inhibitory Factor 1 enhances the production of reactive oxygen species [61] which may activate endothelial cell Ca^2+^ channels [62]. Two observations exclude a role for reactive oxygen species in the control of vascular tone by the mitochondrial ATP synthase: 1) endothelial Ca^2+^ activity is insensitive to global or mitochondrial-targeted scavenging of reactive oxygen species [34]; 2) endothelial vasodilator signaling was abolished by ATP synthase inhibition in the presence of global/targeted reactive oxygen species scavengers. Mitochondrial Ca^2+^ buffering activity is equally unlikely to account for our findings. Indeed, because of the low affinity of the mitochondrial calcium uniporter for Ca^2+^ (kd ~20-30 μM), mitochondria must be closely coupled to Ca^2+^ channels for the organelle to effectively buffer Ca^2+^ release. Ca^2+^ levels sufficient to be captured by mitochondria only occur in the immediate vicinity of active Ca^2+^ sources (distances less than a few hundred nanometers) [63]. In native endothelial cells, mitochondria are mobile organelles that, at any moment in time, are on average a distance of ~1 μm from Ca^2+^ release sites. At such a distance, Ca^2+^ levels are unlikely to rise more than 20 nM above resting values (~100 nM) [64], making the possibility that mitochondria buffer endothelial Ca^2+^ release unlikely.

### 3.3. The complexity of endothelial cell metabolism

Our data indicate that respiratory chain–linked metabolism is necessary for endothelial cell Ca^2+^ signaling and vasodilator activity in multiple vascular beds. This link between mitochondrial metabolism and endothelial function is consistent with observations that polarized mitochondria are required for endothelial Ca^2+^ signaling in intact arteries [34,65,66] and in cell cultures [67–70]. Alongside our investigation of basal, flow-, and ACh-evoked endothelial cell activity, historical findings using various mitochondrial inhibitors suggest that a requirement for mitochondrial ATP may be a general feature of endothelial vasodilator responses, irrespective of agonists, vascular bed, or species [71–75].

While the requirement for mitochondria appears to be a common feature of endothelial cell vasomotor control, mitochondrial ATP production is generally considered to be dispensable for endothelial cell function [76]. However, most studies have been carried out on cultured endothelial cells. The cell culture environment significantly alters endothelial cell metabolism [77,78]. Thus, it is important to understand endothelial energy production in as close to *in vivo* conditions as possible. In this regard, RNA sequencing of freshly-isolated single endothelial cells is increasingly being used, without a cell culture step, to characterize endothelial metabolic signatures [79,80]. Such studies reveal a diversity of metabolic profiles (mitochondrial and glycolytic) amongst endothelial cells of various organs [37,81]. Moreover, the metabolic expression patterns exhibit a remarkable level of plasticity corresponding to distinct physiological functions. For example, angiogenic endothelial cells appear to rely on both oxidative phosphorylation and glycolysis as energy sources [82]. Renal endothelial cells upregulate mitochondrial ATP production to survive water deprivation [83], whilst cardiac endothelial cells upregulate glycolysis in response to ischemia [84]. Our studies used an intact tissue model to demonstrate a key role for mitochondrial metabolism in native endothelial cells in intact arteries, and is supported by a recent transcriptomic analysis demonstrating a preference for oxidative phosphorylation in arterial endothelial cells and a progressive switch to glycolysis along the arteriovenous axis [85]. Thus, as the study of endothelial metabolism moves away from cell culture models, the emerging data suggest that mitochondrial energy production is more important to endothelial cell function than currently appreciated.

### 3.4. Perspective

Endothelial cell function depends on an adequate supply of energy and it is well established that most risk factors for cardiovascular disease (e.g., obesity and aging) impact endothelial cell energy production and lead to vascular dysfunction [17]. Many other non-cardiovascular diseases also involve pathological blood vessel function due to endothelial cell dysfunction. For example, impaired endothelial cell control of blood flow is partly responsible for the vascular problems associated with diabetes, and deficient endothelial barrier function characterizes some types of neurodegeneration. Previously, others have validated the idea of targeting endothelial cell glycolysis as an anti-angiogenic therapy to treat cancer. Our work highlighting that arterial endothelial cells require mitochondrial respiration for the control of vascular tone suggests that aberrant mitochondrial energy production may underlie endothelial dysfunction. As such, therapeutic intervention to promote endothelial cell mitochondrial energy production may be an effective strategy to combat vascular dysfunction in a range of diseases.

## 4. Materials and methods

### 4.1. Animals

All animal care and experimental procedures were conducted in accordance with relevant guidelines and regulations, with ethical approval of the University of Strathclyde Local Ethical Review Panel, and were fully licensed by the UK Home Office regulations (Animals (Scientific Procedures) Act 1986, UK) under Personal and Project License authority. Animal studies are reported in compliance with the ARRIVE guidelines [86].

Experiments were performed on arteries from male or female (8-12 weeks old) Sprague-Dawley rats and female (4-6 weeks old) C57BL6 mice from in house colonies. The animals were housed at The University of Strathclyde Biological Protection Unit. All animals were kept on a 12:12 light/dark cycle (temperature of 21°C ± 2°C, humidity of 45-65%) under standard group housing conditions with unlimited access to water and chow (Rat and Mouse No.1 Maintenance, 801151, Special Diet Services, UK). Animals were kept in RC2F (rats) or M3 (mice) cages (North Kent Plastic, UK) with aspen wood chew sticks and hanging huts for enrichment. Animals were euthanized by cervical dislocation with secondary confirmation via decapitation in accordance with Schedule 1 of the Animals (Scientific Procedures) Act 1986.

Following euthanasia, mesenteric arcades, brains, hearts, or kidneys were removed and transferred to a physiological salt solution (PSS) of the following composition (mM): 125.0 NaCl, 5.4 KCl, 0.4 KH_2_PO_4_, 0.3 NaH_2_PO_4_, 0.4 MgSO_4_, 4.2 NaHCO_3_, 10.0 HEPES, 10.0 glucose, 2.0 sodium pyruvate, 0.5 MgCl_2_, 1.8 CaCl_2_ (adjusted to pH 7.4 with NaOH).

### 4.2. Chemicals

Cal-520/AM (ab171868) and mitoTEMPO (ab144644) were purchased from Abcam (UK). Caged IP_3_ (cag-iso-2-145-100) was purchased from SiChem (Germany). Acetylcholine (A6625), GSK1016790A (G0798), oligomycin (O4876), MSDC-0160 (mitoglitazone, SML1884), phenylephrine (P6126), rotenone (R8875), sodium nitroprusside (S0501), TEMPOL (4-HYDROXY-TEMPO; 176141), and UK5099 (PZ0160) were purchased from Sigma-Aldrich (USA). Pluronic F127 (P3000MP) was purchased from ThermoFisher (UK). Collagenase (Type 2; LS004176) was purchased from Worthington (USA). All other chemicals were obtained from Sigma-Aldrich.

### 4.3. Isolated artery preparation

Either 3^rd^/4^th^ order mesenteric arteries, 1^st^ order posterior cerebral arteries, the septal coronary artery, or interlobular renal arteries were used for measurement of vascular reactivity or endothelial cell Ca^2+^ signalling. All arteries were rapidly dissected, cleaned of fat and adherent tissue and used immediately. Arteries were opened longitudinally, stretched to their *in vivo* length, and mounted on a Sylgard block insert of a custom perfusion chamber using 50 μm diameter tungsten pins. The endothelium was preferentially loaded with the membrane-permeant calcium indicator, Cal-520/AM (5 μM; in DMSO with 0.02% Pluronic F-127), at 37°C for 30 minutes. In a subset of experiments, endothelial cells were also loaded with a membrane-permeant, photolabile form of IP_3_ (caged IP_3_, cIP_3_; 1 μM). In these experiments, cIP_3_ was included in the Ca^2+^-indicator solution.

### 4.4. Endothelial cell sheet isolation

Third and fourth order mesenteric artery segments were enzymatically digested to obtain freshly isolated sheets of endothelial cells (as described previously [34,87]). In brief, vessels were cut open to expose the endothelium, cut into small strips, and incubated in collagenase (Type 2, 2 mg ml^-1^) for 22 minutes at 37°C. The supernatant was then removed and the artery strips were gently washed three times in fresh PSS. Endothelial cell sheets were then dispersed using a wide-bored, fire-polished glass pipette. Endothelial cells sheets were added to a glass-bottomed imaging chamber, for Ca^2+^ imaging, or 8-well chamber slides (μ-slides; Ibidi, Germany), for immunocytochemistry.

### 4.5. Immunocytochemistry

Freshly isolated endothelial cell sheets were allowed to adhere for one hour prior to immunolabelling. Cells were fixed in 4% paraformaldehyde (Agar Scientific, UK) in phosphate buffered saline (PBS) for 20 minutes at room temperature. Cells were then washed three times in glycine solution (0.1 M), three times in PBS, and then permeabilized with Triton-X100 (0.2% in PBS) for 30 minutes. Cells were again washed three times in PBS, three times in an antibody wash solution (150 mM NaCl, 15 mM Na_3_C_6_H_5_O_7_, 0.05% Triton-X100 in ddH_2_0), and incubated for 1 hour with blocking solution (5% donkey serum in antibody wash solution) at room temperature. All individual wash steps were 5 minutes in duration. Cells were then incubated overnight at 4°C with goat anti-mouse platelet endothelial cell adhesion molecule (CD31/PECAM-1) primary antibody (R&D Systems cat. No. AF3628) diluted in antibody buffer (1:1000; 150 mM NaCl, 15 mM Na_3_C_6_H_5_O_7_, 2% donkey serum, 1% BSA, 0.05% Triton X-100, 0.02% sodium azide in ddH_2_0). Following primary antibody incubation, cells were washed three times in antibody wash solution, and incubated for 1 hour at room temperature with a fluorescent secondary antibody conjugated with Alexa Fluor 488 (donkey anti-goat, 1:1000; A-11055) in antibody buffer. Cells were then washed three times in antibody wash solution, incubated with the nuclear stain, 4’,6-diamidino-2-phenylindole (DAPI; 4 nM), for 5 minutes, and finally washed three more times in PBS prior to imaging. Fluorescence images of mesenteric artery endothelial cell sheets were acquired using an inverted fluorescence microscope (TE-300; Nikon, Japan) equipped with a 100X, 1.4 numerical aperture oil immersion objective, 400/460 LED illumination (CoolLED, UK), and an iXon 888 (Andor, UK) electron multiplying CCD camera. Images were acquired using μManager software [88].

### 4.6. Assessment of vascular reactivity

Vascular reactivity was assessed in isolated mesenteric arteries mounted en face [28–30]. Arteries were visualized at 5 Hz using an inverted fluorescence microscope (TE2000; Nikon, Japan) equipped with a 20X, 0.75 numerical aperture objective, 460 nm LED illumination (CoolLED, UK), and an iXon 888 (Andor, UK) electron multiplying CCD camera. The resulting 666 x 666 μm field of view allowed quantification of vascular reactivity in opened arteries using VasoTracker edge-detection algorithms [89]. All arteries used for experimentation had a luminal diameter ~150 μm, and were perfused with PSS at a rate of 1.5 ml min^-1^. PE-induced constriction was expressed as the percentage reduction from resting diameter. ACh-evoked vasodilation was expressed as a percentage of maximal relaxation (constricted diameter to resting diameter).

Arteries were partially constricted with phenylephrine added to the perfusate (to ~80% of resting diameter; 20% contraction; ~2 μM phenylephrine). This level of constriction enables the vessels to either dilate or constrict further under experimental pharmacological studies [28]. Endothelial function was then assessed by the vasodilator response to ACh (10 μM). Under these control conditions, all arteries constricted to PE, exhibited >80% relaxation to ACh, and were included in subsequent analysis. Following washout, vasoconstrictor/vasodilator responses were examined once more. Underlying mechanisms of ACh-evoked vasodilation were then examined by adding additional pharmacological agents (as described in the text) to the perfusing bath solution. These experiments allowed us to test whether the drugs were able to reverse ACh-evoked vasodilation. In a subset of experiments, arteries were incubated with pharmacological agents before the second test of vascular reactivity. These experiments allowed the effects of the drugs on PE-evoked constriction and ACh-evoked vasodilation to be examined. Sodium nitroprusside (100 μM) was added to the perfusate at the end of each experiment to confirm endothelium-independent vasodilation.

### 4.7. Endothelial cell calcium imaging

Intact mesenteric, cerebral, coronary, and renal artery endothelial cell Ca^2+^ activity was recorded at 10 Hz using the same microscope system used for assessment of vascular reactivity, but using a 40x, 1.3 numerical aperture oil immersion objective. The resulting 333 x 333 μm field of view was used to visualize large endothelial cell networks (~250 cells). A paired experimental was used to examine basal (unstimulated/spontaneous), flow-evoked, and ACh-evoked Ca^2+^ activity before and after treatment with various pharmacological agents (40-minute incubation), as described in the text. Basal activity (2 minutes) was examined in the absence of flow or pharmacological agents. Flow-evoked activity was examined during/after the initiation flow at a rate of 1.5 ml min^-1^ (2-minute recording; 30 second baseline). ACh-evoked activity was examined by adding the drug to the perfusate (2-minute recording; 30 second baseline). In a separate series of experiments, TRPV4-mediated intact artery endothelial Ca^2+^ responses were evoked by the specific TRPV4 channel agonist, GSK1016790A. In another, we examined Ca^2+^ responses evoked by the photolysis of caged IP_3_, using a computer-controlled tripled neodymium: yttrium aluminum garnet (Nd:Yag; wavelength 355 nm) laser (Rapp Optoelektronic, Germany) attached directly to the TE2000 microscope system [35]. In yet another series of experiments, we examined ACh-evoked activity in isolated sheets of mesenteric artery endothelial cells. Again, ACh was added to the perfusate and a paired experimental design was employed to examine the effects of pharmacological manipulation on endothelial Ca^2+^ signaling. All images were acquired using μManager software [88].

### 4.8. Analysis of calcium activity

Single-cell endothelial Ca^2+^ activity was assessed as previously described [30,35]. In brief, we used automated algorithms to extract fluorescence intensity as a function of time from circular regions of interest (6.5 μm diameter) centered on each cell in our images. Fluorescence signals were then smoothed using a Savitzsly-Golay (21 point, 3^rd^-order) filter, expressed as fractional changes in fluorescence (F/F_0_) from baseline (F_0_). The baseline was automatically determined by averaging the fluorescence intensity of the 100-frame portion of each trace that exhibited the least noise. We then calculated the discrete derivative (d(F/F_0_)/dt) of each Ca^2+^ signal, and used a peak-detection algorithm to identify increases in fluorescence intensity rising at least 10 standard deviations above baseline noise. Endothelial Ca^2+^ activity was quantified using the number (or percentage) of cells exhibiting spiking activity, and the amplitude of these Ca^2+^ spikes (ΔF/F_0_).

### 4.9. Statistical analysis

For all experiments, the reported n represents the number of biological replicates (number of animals). Summary data are presented in text as mean ± standard error of the mean (SEM), and graphically as mean ± SEM (time-course data) or individual data points with the mean ± SEM indicated. Paired data in plots are indicated by the shape and colour of the plotted points. Data were analyzed using paired t tests, independent 2-sample t tests (with Welch’s correction as appropriate), repeated measures one-way ANOVA with Tukey’s or Dunnett’s test for multiple comparisons, as appropriate as in the respective figure legend. All statistical tests were two-sided. A p value of < 0.05 was considered statistically significant.

## Data Availability

All data underpinning this study is available from the authors upon reasonable request.

## Acknowledgements

This work was funded by the Wellcome Trust (204682/Z/16/Z; 202924/Z/16/Z) and the British Heart Foundation (RG/F/20/110007; PG/16/54/32230; PG/20/9/34859), whose support is gratefully acknowledged.

## Video Legends

**Video 1: Inhibition of the mitochondrial ATP synthase reverses acetylcholine-evoked vasodilation.**

Representative videos, without (left) and with (right) VasoTracker tracking results overlaid, showing the effect of oligomycin on the vasodilation response of a third order mesenteric artery. Arteries were splayed open, pinned flat, before endothelial cells were labelled with the fluorescent indicator, Cal-520/AM (5 μM), and visualized using wide field single-photon microscopy (16X objective, NA = 0.8, ~0.69 mm^2^ field of view, 0.66 μm^2^ projected pixel size). The width of artery segments (unfolded circumference) was converted to the equivalent diameter to illustrate the time course of the mechanical response (bottom). At the plateau phase of contractions evoked by phenylephrine (PE, titrated to achieve 20% contraction), acetylcholine (ACh, 10 μM) caused relaxation of the preparations. Subsequent addition of the mitochondrial ATPase inhibitor, oligomycin (2.4 μM), reversed this relaxation and increased contraction to levels greater than that induced by PE alone. Videos show the central section (832 μm × 832 μm field of view) of an ~ 3 mm long blood vessel at pixel size = 0.66 μm^2^/pixel. Data also displayed in Figure 1.

**Video 2: Spontaneous endothelial calcium signaling requires mitochondrial ATP production.** Representative videos of basal (spontaneous) endothelial Ca^2+^ activity before (left) and after (right) incubation with the mitochondrial ATPase inhibitor, oligomycin (2.4 μM). Arteries were splayed open and pinned flat before endothelial cells were preferentially labelled with the fluorescent Ca^2+^ indicator, Cal-520/AM (5 μM), and visualized using high-resolution wide field single-photon microscopy (40X objective, NA = 1.3, 333 μm × 333 μm field of view, 0.11 μm^2^ projected pixel size). Ca^2+^ imaging data is displayed as fractional change in fluorescence movies (F/F_0_). Ca^2+^ traces are shown for 5 randomly selected cells, and are illustrated as a heatmap for all 197 cells contained within the field of view. Under control conditions (left), quiescent endothelium exhibits substantial spontaneous Ca^2+^ activity. After incubation with oligomycin (right), quiescent endothelium exhibit little spontaneous Ca^2+^ activity. The dataset in the movie is shown in summarized data (Figure 6D) as blue-outlined squares.

**Video 3: Flow-evoked endothelial calcium signaling requires mitochondrial ATP production.** Representative videos of flow-evoked (1.5 ml min^-1^) endothelial Ca^2+^ activity before (left) and after (right) incubation with the mitochondrial ATPase inhibitor, oligomycin (2.4 μM). Arteries were splayed open and pinned flat before endothelial cells were preferentially labelled with the fluorescent Ca^2+^ indicator, Cal-520/AM (5 μM), and visualized using high-resolution wide field single-photon microscopy (40X objective, NA = 1.3, 333 μm × 333 μm field of view, 0.11 μm^2^ projected pixel size). Ca^2+^ imaging data is displayed as fractional change in fluorescence movies (F/F_0_). Ca^2+^ traces are shown for 5 randomly selected cells, and are illustrated as a heatmap for all 197 cells contained within the field of view. Prior to oligomycin treatment (left), flow of physiological saline amplifies spontaneous endothelial Ca^2+^ activity. After incubation with oligomycin (right), the endothelium exhibits little spontaneous Ca^2+^ activity and flow is without effect. The dataset in the movie is summarized in Figure 6D, and shown as blue-outlined squares.

**Video 4: Acetylcholine-evoked endothelial calcium signaling requires mitochondrial ATP production.** Representative videos of acetylcholine (ACh)-evoked (10 μM) endothelial Ca^2+^ activity before (left) and after (right) incubation with the mitochondrial ATPase inhibitor, oligomycin (2.4 μM). Arteries were splayed open and pinned flat before endothelial cells were preferentially labelled with the fluorescent Ca^2+^ indicator, Cal-520/AM (5 μM), and visualized using high-resolution wide field single-photon microscopy (40X objective, NA = 1.3, 333 μm × 333 μm field of view, 0.11 μm^2^ projected pixel size). Ca^2+^ imaging data is displayed as fractional change in fluorescence movies (F/F_0_). Ca^2+^ traces are shown for 5 randomly selected cells, and are illustrated as a heatmap for all 197 cells contained within the field of view. Prior to oligomycin treatment (left), the addition of ACh to the flowing physiological saline evokes a large, global increase in endothelial Ca^2+^ levels. After incubation with oligomycin (right), the endothelium exhibits little Ca^2+^ activity despite ongoing flow, and the response to ACh is muted. The dataset in the movie is summarized in Figure 6D, and shown as blue-outlined squares.

**Video 5: The endothelial requirement for mitochondrial ATP is preserved in freshly isolated cells.** Representative videos of acetylcholine (ACh)-evoked (10 μM) Ca^2+^ activity in an isolated sheet of endothelial cells before and after incubation with the mitochondrial ATPase inhibitor, oligomycin (2.4 μM). Endothelial cells were enzymatically dissociated from mesenteric arteries and labelled with the fluorescent Ca^2+^ indicator, Cal-520/AM (5 μM), and visualized using high-resolution wide field single-photon microscopy (40X objective, NA = 1.3, 333 μm × 333 μm field of view, 0.11 μm^2^ projected pixel size). Ca^2+^ imaging data is displayed as pseudocolored fractional change in fluorescence (F/F_0_). Ca^2+^ traces were extracted automatically from all cells and traces are shown: for 5 randomly selected cells, and as a heatmap for all 210 cells contained within the visualized sheet. Prior to oligomycin treatment (left), the addition of ACh to the flowing physiological saline evokes a large, global increase in endothelial Ca^2+^ levels. After incubation with oligomycin (right), the response to ACh is muted. The dataset in the movie is summarized in Figure 6.

**Video 6: Directly-evoked IP_3_-mediated endothelial calcium signalling requires mitochondrial ATP synthesis.** Representative videos of IP_3_-evoked endothelial Ca^2+^ activity before (left) and after (right) incubation with the mitochondrial ATPase inhibitor, oligomycin (2.4 μM). Arteries were splayed open and pinned flat before endothelial cells were preferentially labelled with a fluorescent Ca^2+^ indicator (Cal-520/AM, 5 μM) and caged IP_3_, and visualized using high-resolution wide-field single-photon microscopy (40X objective, NA = 1.3, 333 μm × 333 μm field of view, 0.11 μm^2^ projected pixel size). Ca^2+^ imaging data is displayed as fractional changes in fluorescence movies (F/F_0_). Ca^2+^ activity was evoked by laser targetted photorelease of IP_3_ in four regions (indicated by coloured circles). Average traces for each region are shown. Oligomycin inhibits Ca^2+^ signals evoked by laser-targeted photorelease of IP_3_.

